# The Interplay Between Sketching and Graph Generation Algorithms in Identifying Biologically Cohesive Cell-Populations in Single-Cell Data

**DOI:** 10.1101/2023.09.15.557825

**Authors:** Emma Crawford, Alec Plotkin, Jolene Ranek, Natalie Stanley

## Abstract

High-throughput single-cell immune profiling technologies, such as mass cytometry (CyTOF) and single-cell RNA sequencing measure the expression of multiple proteins or genes across many individual cells within a profiled sample. As it is often of interest to identify particular *clusters* or cell-populations driving clinical phenotypes or experimental outcomes, there is a critical need to develop automated bioinformatics approaches that can handle a large number of profiled cells. For analyzing multi-sample single-cell datasets at scale, the datasets are usually encoded as a graph, where nodes represent cells and edges imply significant between-cell similarity. As multi-sample single-cell experiments can readily result in millions of profiled cells, the construction and analysis of a graph becomes computationally prohibitive and often requires reducing the dataset size through downsampling as a pre-processing step. Here, we explore the interplay between sketching, or downsampling approaches, and the way in which the graph is constructed on the sketched data for ultimately identifying biologically-meaningful cell-populations. Our results suggest that combining a principled sketching approach with a simple *k*-nearest neighbor graph representation of the data can identify meaningful subsets of cells as robustly as, and sometimes better than, more sophisticated graph generation approaches. This reveals that the practical concern of downsampling or sketching a limited number of cells is a more critical pre-processing step than how the graph representation is constructed.

## 1 Introduction

Advancements in high-throughput immune profiling techniques, such as mass cytometry (CyTOF) [24] and single-cell RNA sequencing (scRNA-seq) [19] have enabled the comprehensive characterization of diverse immune cell-types present in a single blood or tissue sample, through the simultaneous measurement of several proteins or genes, respectively. Such technologies are often used for clinical phenotyping in a variety of settings, including characterizing surgical recovery [18], predicting deleterious pregnancy outcomes [25], or predicting immunotherapy resistance [13]. For application in such settings, single-cell data are often collected from multiple patient samples, which can collectively generate feature vectors for millions of cells. Each cell is represented as a feature vector in *d* dimensional space, where *d* is the number of genes or proteins measured per cell. A primary goal of studying these data is to identify particular immune cell-types or *populations* that correlate with a particular clinical outcome or experimental condition [5, 21, 22, 29, 23].

Two complementary approaches are commonly used to identify cell-populations [5, 21] or prototypical cells [22, 23, 29] driving the clinical phenotype or experimental condition of interest. Under the cell-population level approach, cells are often clustered according to the *d* measured gene or protein features [5, 21, 4, 6]. Features are then typically engineered on a per-population (cluster) level and ultimately used for downstream clinical outcome prediction tasks. Alternatively, identifying outcome-specific prototypical cells requires scoring individual cells based on the extent to which they and their nearest neighbors are associated with an outcome of interest. The commonality between these two paradigms is that they rely on constructing a graph-based representation of the data. Under these represen-tations, nodes represent individual cells, and an edge between cells implies that they are *phenotypically similar*, or have a small distance between them in the *d*-dimensional feature space. Once the graph is constructed, cell-populations are identified through graph-based clustering [6] or, alternatively, prototypical cells are scored using a graph signal processing approach [22] or based on general properties of cellular neighborhoods [29, 23].

While these methods are useful for identifying immunological correlates with outcomes of interest, they suffer from a practical limitation: they often fail to accommodate the potentially millions of cells that are profiled in a multi-sample single-cell experiment [28]. In particular, the two bottlenecks in the analysis of a large number of cells are (1) the construction of the graph and (2) subsequent downstream tasks like graph partitioning [15, 2] and signal filtering [17]. Moreover, it is unclear how sparse the graph must be in order to extract meaningful cell-populations or to compute accurate per-cell outcome-associated scores. To aid in the scalability of graph construction, researchers often implement a downsampling or *sketching* approach to select a limited subset of representative cells for downstream analysis [27, 14, 10]. It is common to use a *k*-nearest neighbor graph to construct a graph representation of the data, [6, 23, 22], but recent advances in graph structure learning have also suggested more principled approaches for constructing well-structured graphs [30]. Both the sketching and graph construction approaches have strong potential to impact downstream analyses independently (see Sections 1.1 and 1.2 for a detailed description of the nuances of these approaches), but their interplay has not been explored or characterized. In this work, we systematically explore the implications of sketching and graph construction on downstream analysis tasks and specifically find that leveraging a smart (i.e. not uniformly random) sketching method significantly aids in performance.

### 1.1 Overview of Sketching Approaches

Single-cell datasets are information-rich and contain diverse cell-populations with different geometries and relative abundances. When selecting a limited number of cells through a downsampling or sketching approach, it is critical to select cells that are in line with the underlying geometry and to not artificially oversample cells from dense, well-populated cell types. A range of sketching approaches for single-cell data have already been proposed, including Kernel Herding[27], Geometric Sketching[14], and Hopper[10], that seek to efficiently capture salient information about the data using a small number of cells. The Geometric Sketching and Hopper methods are most focused on approximating the original data geometry as closely as possible, whereas Kernel Herding Sketching is most concerned with maintaining the distribution of cells in the original data. All of these approaches have strong theoretical background to support them, which we will explore in detail here.

Kernel Herding (KH) sketching is based on the Radial Basis Function kernel popular in machine learning [27]. The Radial Basis Function kernel is part of a special class of kernels that enable a function called the *kernel mean embedding* to uniquely characterize distributions. Because of this property, if two distributions have close kernel mean embeddings, the distributions themselves must be close. Thus, in Kernel Herding Sketching, the objective function to minimize is the distance between the kernel mean embedding of the original data and that of the subsampled data. Because the RBF kernel is expensive to compute explicitly, an approximation is constructed using random Fourier frequencies. Kernel Herding has been shown to maintain the relative frequencies of cell types as a uniformly random sample would, but it is significantly better at maintaining the structural properties of the original data geometry (as quantified by the singular values of the feature matrix). However, because the distribution of cell types is close to that of the original data, it is possible to miss rare cell types in the subsample if the sample size is too small.

Geometric Sketching is a method developed for analyzing single-cell data that specifically aims to oversample rare cell types [14]. This is accomplished by volume sampling, a method of approximating the geometry of the data by “covering” it with high dimensional cubes. These cubes are placed so that all of the data points are contained within one of these cubes and each cube is nonempty. Then, the same number of points are chosen from each cube to be in the sample. For common cell types, the ratio of volume they take up to total volume of the feature space is low, as they are generally densely packed in the space. Thus, despite their high relative frequency by count, they end up getting undersampled in this method. Conversely, the rare cell types are spread out in the feature space, so they get oversampled. Oversampling rare cell types is helpful when trying to understand the entirety of the data geometry or analyzing cell-type heterogeneity; however, this oversampling also leads to a distribution of cells that does not match that of the original data. This means that analysis of the subsample can’t be as easily extrapolated to the full dataset.

Similarly to Geometric Sketching, the Hopper Sketching method aims to approximate the high-dimensional data geometry [10]. This is accomplished by finding a subsample of the data so that the Haussdorff distance from the original data to the subsampled data is small. That is, if *X* is the original data, the goal is to find a subset *S* of that data so that

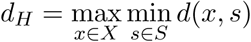

is minimized. This is accomplished by iteratively finding the point in the original data that is farthest from the subsample. This point is then added to the subsample and the process is repeated with the new subsample. Given that the goal is approximating the data geometry, the Hopper Sketching method ends up oversampling rare cell types in a similar manner to Geometric Sketching. Hopper Sketching has a pre-partitioned version of their algorithm for large datasets, which we will detail in Section 2.2. This pre-partitioning makes Hopper the fastest of the sketching methods that we studied.

Finally, IID sampling creates a discrete uniform probability distribution over the *n* cells, so each cell has a 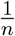 probability of being included in the sample. Then, the desired number of cells are chosen from this distribution without replacement. This method is extremely time-efficient and maintains relative frequencies of cell types from the original data, but it does not use any information about the data; therefore, it could miss rare cell types or important parts of the data geometry.

### 1.2 Overview of Graph Generation Approaches

Creating a cell-similarity graph from single-cell data provides a useful data structure for downstream analysis tasks, such as discovering subsets of highly similar cells [6]. Here, nodes of the graph represent cells, and edges between cells imply that a given pair of cells are sufficiently similar in the measured feature space. A simple and widely used approach for creating a graph representation of single-cell data is to construct a *k*-nearest neighbor, or *k*-NN, graph.

Briefly, to generate this graph from data with *d* protein or gene measurements, we view each cell as a point in *d*-dimensional space. If a cell has measurements *m*_1_, …, *m*_*d*_, then the coordinates of that point are exactly (*m*_1_, …, *m*_*d*_). Then for each cell, we place edges connecting the cell to each of the *k* closest nodes to it based on a predetermined metric (Euclidean distance or cosine distance, e.g.). The weight of an edge between cells *c*_1_ and *c*_2_ is defined as 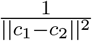, where || *·* || denotes the aforementioned distance metric, so that cells that are close together have strong edges between them. Using *k*-NN graphs is popular because they are intuitively simple and create highly structured, *sparse* (i.e. with few edges) graph representations; however, it is often difficult to determine the appropriate *k* for each dataset, as the optimal value of this parameter is likely to vary depending on the data geometry. When *k* is small, the analysis is fast due to the sparsity of the graph, but the resulting graph is not well-connected and critical information about the underlying manifold structure may be lost. When *k* is too large, spurious or redundant edges are added, which inserts noise into the analysis. High graph density could also significantly slow down the downstream analysis of the graph.

More sophisticated graph construction algorithms leverage theory from the field of *graph signal processing* [7, 11, 1, 3, 9, 8]. Rather than viewing the *d* protein or gene measurements in a dataset as dimensional coordinates, these algorithms view them as functions or *signals*, for example as *g*_1_, …, *g*_*d*_, so that *g*_*i*_(*j*) is the *i*^*th*^ measurement of the *j*^*th*^ cell. The goal of this class of algorithms is to therefore create a graph representation of the data such that these signals vary smoothly over the graph [7]. Because there are many smoothness constraints in this problem formulation, the algorithms generally pose the constraints as a convex optimization problem, as in [1, 9, 8]. Solving these can be computationally prohibitive, especially on large datasets. Scalable versions of these algorithms generally focus on the eigenvalues of the graph Laplacian, since advancements in spectral methods have made such computations much more efficient [26, 32, 12]. One example of this is the GRASPEL algorithm[30]. The graph formation starts with a 2-NN graph as its base, then iteratively adds the most “spectrally significant” edges until the spectrum has stabilized up to a predetermined tolerance. The spectral significance of an edge is a measure of how much the eigenvalues of the graph Laplacian would differ before and after adding the edge. The spectrally significant edge candidates are found by first sorting the nodes by the eigen-vector of the second smallest eigenvalue of the current graph Laplacian (one property of graph Laplacians is that the first eigenvalue is always zero). This vector is often called the *Fiedler vector*. Then, edges are placed between nodes at the top and nodes at the bottom of this vector. Edges between these nodes are likely to be spectrally significant because the Fiedler vector encodes a suggested partition of the data into two subsets: those with negative entries in the Fiedler vector and those with positive entries. Thus, connecting nodes from the top and the bottom is connecting previously very disconnected regions of the graph. As more disconnected regions of the data become connected via this process, the original data geometry becomes more well-approximated, and so the graph Laplacian spectrum (which encodes the data geometry) also stabilizes. Given that the spectrum of a graph generally stabilizes very quickly when adding edges this way, the algorithm produces an ultra-sparse graph (averaging less than two edges per node), which makes downstream tasks such as clustering much faster.

### 1.3 Summary of Contributions

In this paper, we generate graphs using combinations of sketching and graph generation methods by first downsampling the data and then creating a graph representation of that subsample. We evaluate our results on the ability to identify quality cellular subsets and in a semi-supervised learning task for partially-labeled data cell population identification and semi-supervised learning tasks. Key insights gleaned from our work can further be summarized as follows:

- Sufficiently dense graphs generated from sketches that maintain relative frequencies of the original data tend to produce graph structures which reveal high-quality biologically cohesive cellular subsets.
- In semi-supervised learning tasks such as label propagation, where the objective is to use a certain proportion of labeled nodes to label the unlabeled nodes, the sketching method has much more of an impact on performance than the graph generation method. Specifically, samples generated through Kernel Herding Sketching and Geometric Sketching are ideal.
- While a particular density is required in the graph to identify salient cellular subsets, the usefulness of *k*-NN graphs diminishes for *k >* 8.

## 2 Methods

In this work, we systematically explored all possible combinations of five graph generation methods and four sketching methods on the ultimate discovery of biologically cohesive cell-populations.

### 2.1 Notation and Definitions

#### Basic Data Structure

Single-cell data is typically encoded in a standard data matrix, *F* with *d* columns of protein or gene measurements and *n* rows of cells. Thus, the (*i, j*) entry of the matrix *F*_*i,j*_ has the *j*^*th*^ measurement from the *i*^*th*^ cell. The cells also often come with discrete *labels* describing particular cellular properties, such as the day the cell was sampled, the disease status of the patient that the cell was taken from, or the cell type.

#### Notion of Distance for Graph Construction

In graph generation, we think of these measurements as corresponding to coordinates in a high-dimensional *feature space*. Because of this view, we talk about the rows of the feature matrix as indicating the position of the cells. For conciseness, instead of using the row notation *F*_*i*,:_, we will refer to to the position vector of the *i*^*th*^ cell as **x**_*i*_. An important part of graph generation is calculating the distance between points in this space; we will denote the Euclidean distance between the *i*^*th*^ and *j*^*th*^ cells as ||**x**_*i*_ − **x**_*j*_||.

#### Reducing Dimension Prior to Clustering

When there are many measurements in the dataset, we perform *principal component analysis* to reduce the dimension of the feature matrix while losing minimal information. Biological data can generally be well-represented using only around 30-50 dimensions; principal component analysis allows us to generate a new feature matrix with the same number of rows (cells), but only 30-50 columns. This helps to get rid of both noise and redundancy in the data, and it speeds up computations.

#### Creating and Mining the Similarity Graph

Finally, a *cell similarity graph* is a set of *nodes* connected by *edges* where each node represents a cell and edges encode similarity between them. If node *i* and node *j* are connected by an edge, we denote this as *e*_*ij*_ ∈ *E*, and we call *E* the *edgelist*. Cell-similarity graphs are generally *weighted, undirected*, and *simple*. A weighted graph is one where each edge has a value assigned to it; large edge weights indicate strong connections between nodes. In undirected graphs *e*_*ij*_ ∈ *E* if and only if *e*_*ji*_ ∈ *E*. Finally, simple graphs have no edges of the form *e*_*ii*_; that is, there are no edges from one node to itself. For computational purposes, we store the edgelist information in an *n × n* matrix *A*, called the *adjacency matrix*. If there is an edge between nodes *i* and *j* with weight *w*, then *A*_*ij*_ = *w* (and *A*_*ji*_ = *w*, since the graph is undirected). The diagonal entries of this matrix are all zeros, since the graph is simple. From this adjacency matrix, we can form the *graph Laplacian* matrix. The off-diagonal entries of the graph Laplacian are exactly those of the adjacency matrix (so *L*_*ij*_ = *A*_*ij*_ whenever *i* ≠ *j*), but the diagonal entries are defined as

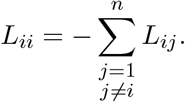

This construction forces the first eigenvector of this matrix to be **1**, an *n ×* 1 vector consisting of all 1s, corresponding to the eigenvalue 0. Since this property is shared across all graph Laplacians, the *Fiedler vector*, or an eigenvector corresponding to the second smallest eigenvalue, is the most important vector in discussion of the graph Laplacian spectrum.

### 2.2 Implementation Details for Sketching Methods

For each sketching method, we generated samples ranging in size from 100 cells to 700 cells, in increments of 100. To balance information and readability, we present only the results from 300-, 500-, and 700-node samples for Figures 2 and 3. The scatter plots in Figures 4 and 5, along with the boxplots in Figure 6, reflect results from samples with 700 nodes. We obtained 30 samples of each size via each sketching method.

**Figure 1:**
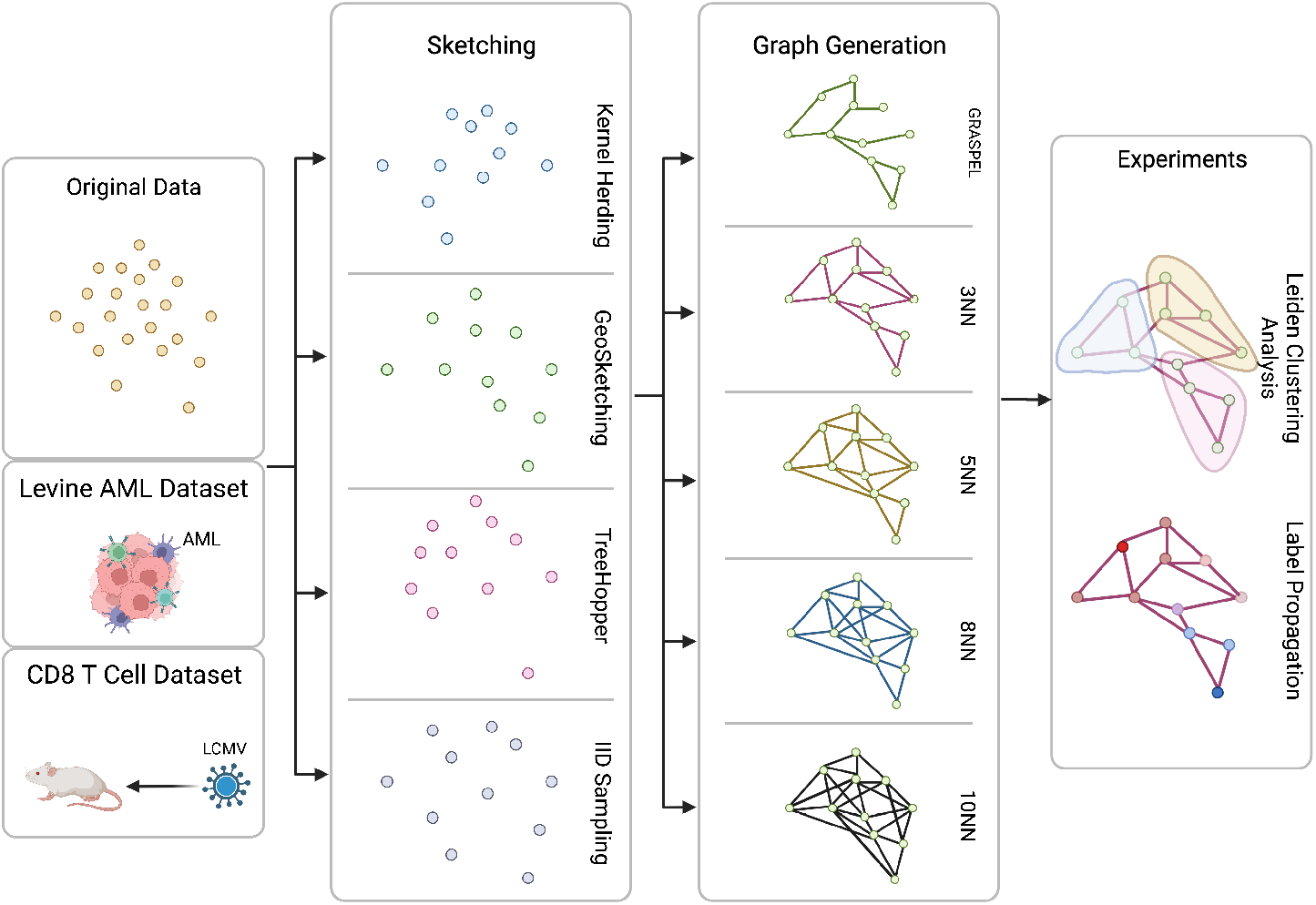
We evaluate the interplay between sketching and graph generation methods on the discovery of biologically cohesive cell-populations in single-cell datasets. Two single-cell datasets, four sketching methods, and five graph generation approaches are used to objectively and quantitatively evaluate the quality of cell-populations uncovered in the dataset. Created with BioRender.com.

**Figure 2:**
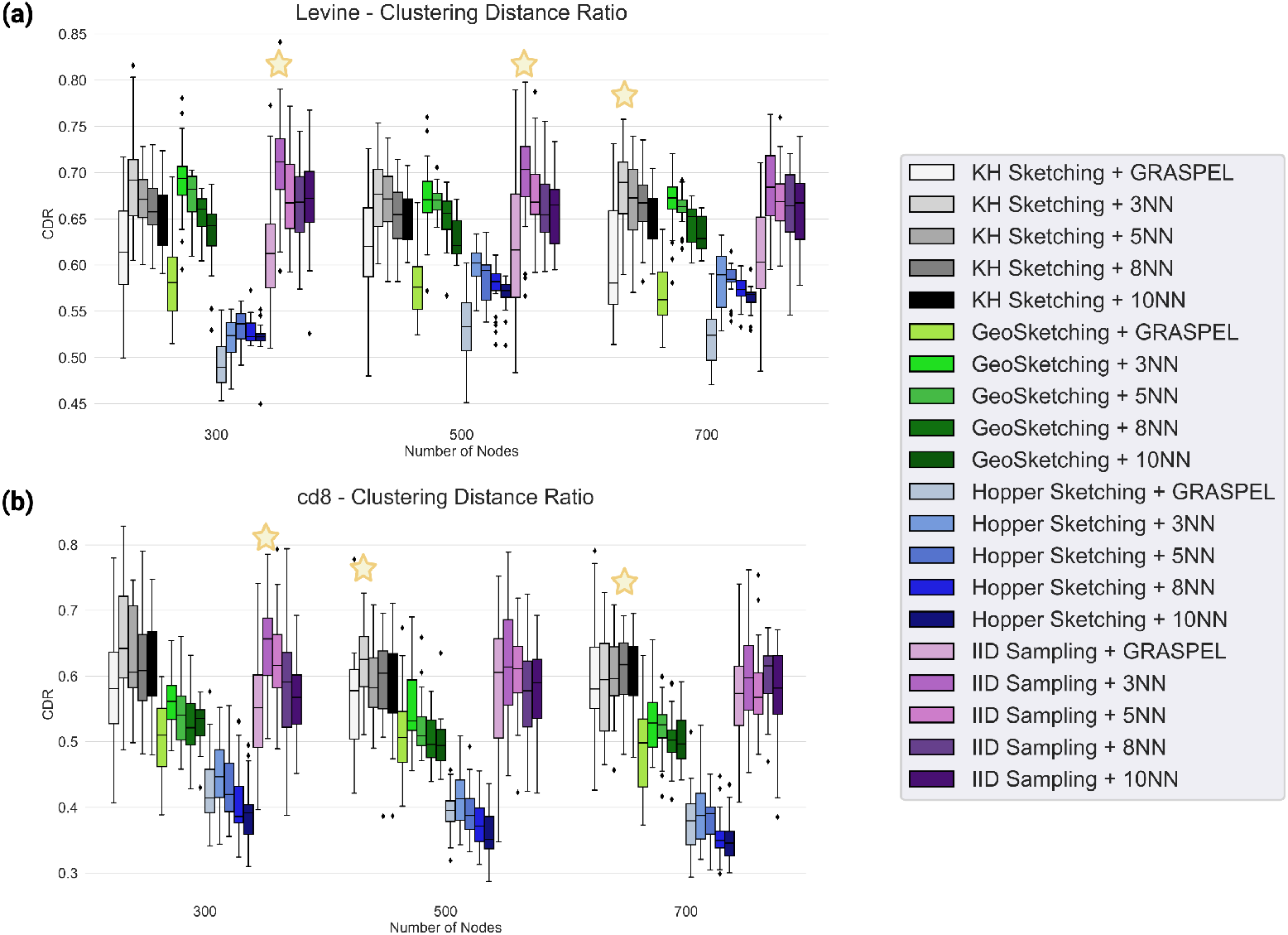
A Clustering Distance Ratio (CDR) was computed to quantify how well-separated clusters are under each combination of sketching and graph generation approach. Boxplots visualize the distribution over 30 sketches in the the Levine dataset (a) and in the T-cell dataset (b). The star indicates the highest median CDR for each number of nodes. In both datasets, the graph density is overall negatively correlated to the CDR, with the exception of the poor performance of the ultra-sparse GRASPEL graph.

**Figure 3:**
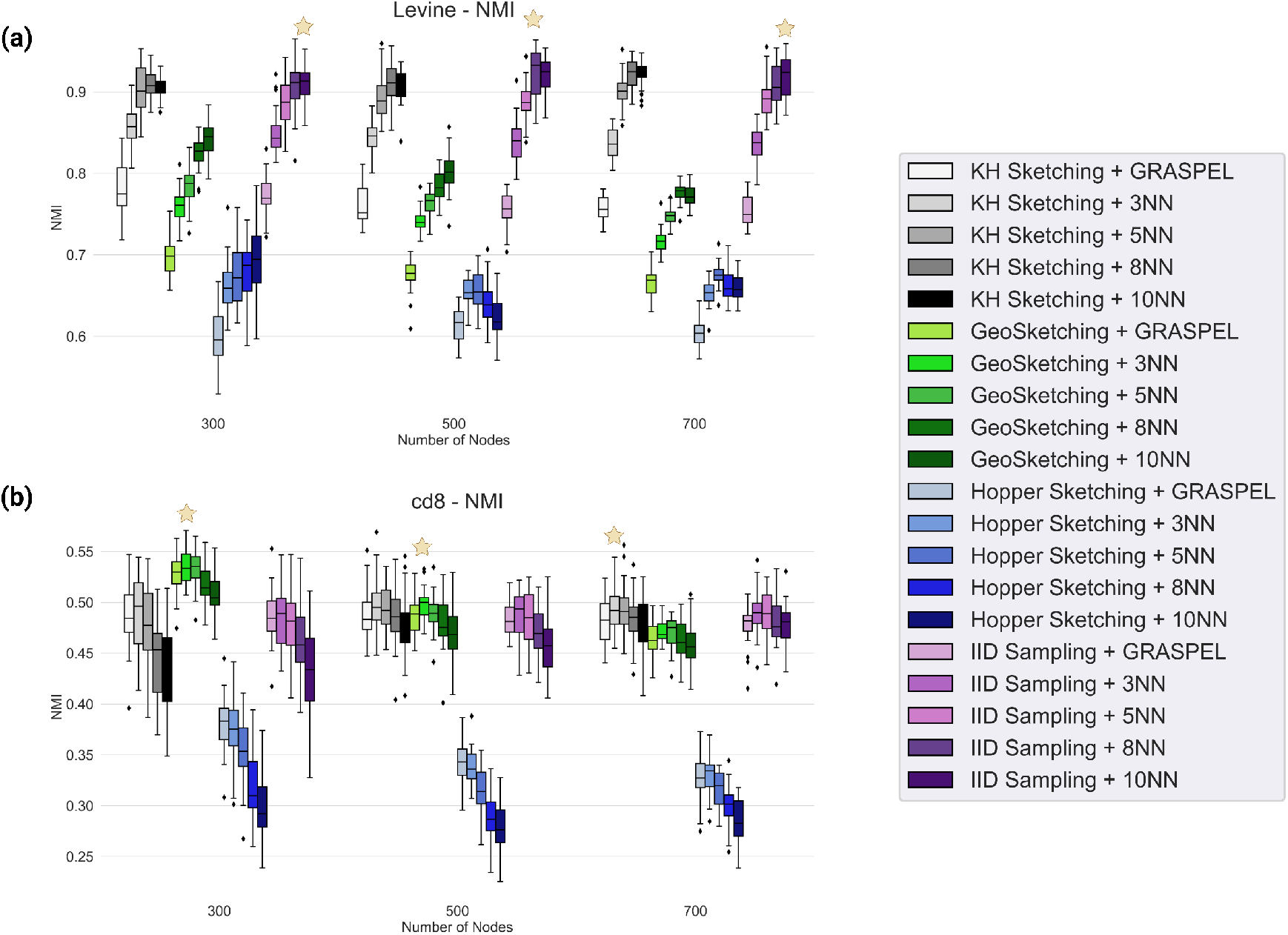
The quality of cluster assignments in relation to the true labels of the cells were quantified with NMI. Boxplots visualize the distribution of NMIs over 30 sketches with each method and are shown for (a) the Levine dataset and (b) the CD8 T-cell dataset. The star indicates the highest median NMI for each number of nodes. As more edges are added to the graph, the NMI increases. IID sampling with an 8-NN graph performs the best on the Levine dataset, with Kernel Herding and an 8-NN or 10-NN graph performing similarly well for all numbers of nodes.

**Figure 4:**
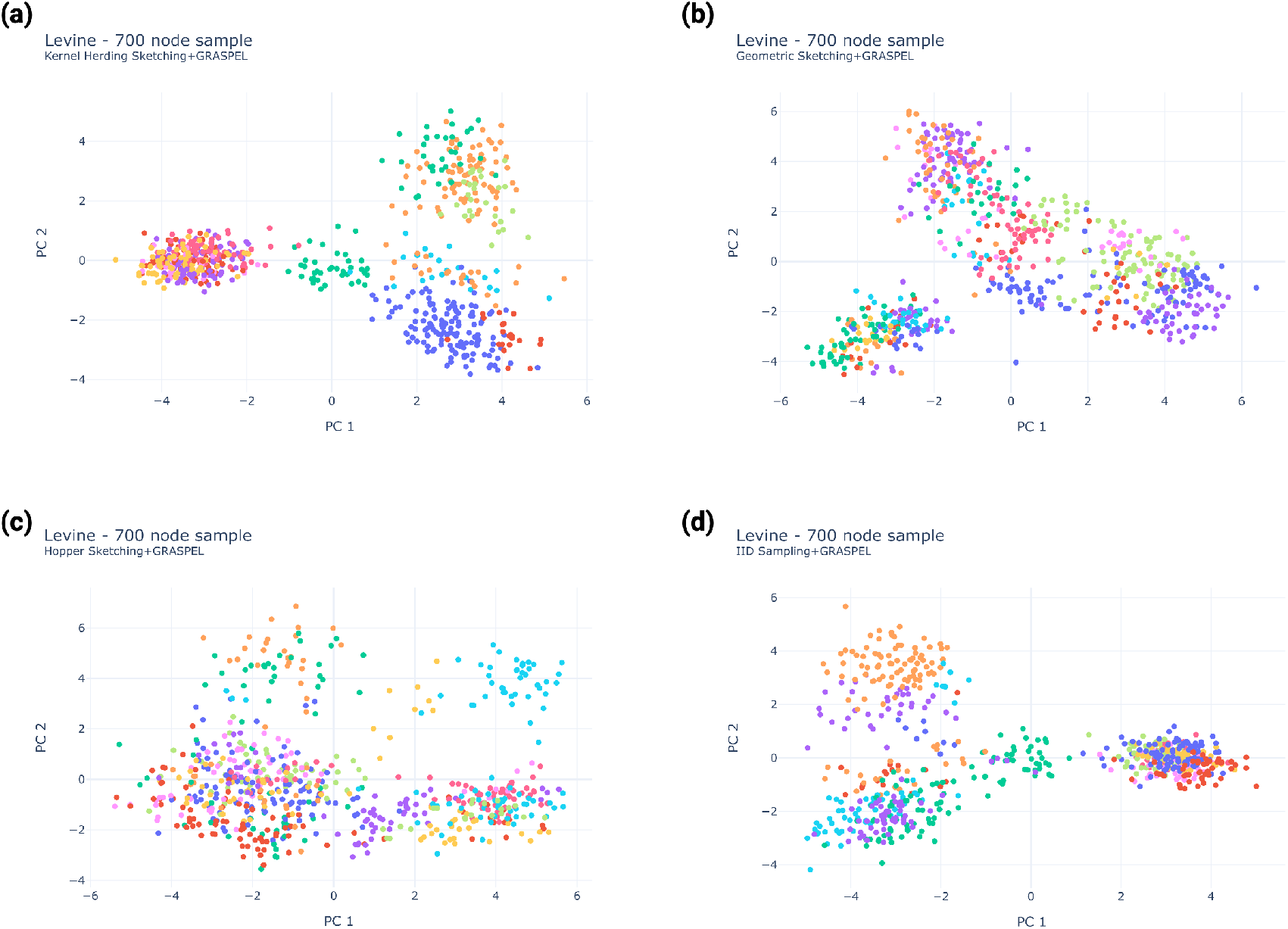
Samples of 700 cells from the Levine AML dataset projected in two dimensions with PCA and colored according to Leiden clustering assignments. All sketching methods are shown here and include (a) Kernel Herding Sketching, (b) GeoSketching, (c) Hopper Sketching, and (d) IID sampling. GRASPEL was used as the graph generation method in all of these plots. The IID sample and Kernel Herding sketch have very similar geometry, explaining their close performance with respect to the clustering distance ratio and the NMI metrics. The Hopper data has lots of overlapping and spread-out clusters, making distinguishing cell types difficult.

**Figure 5:**
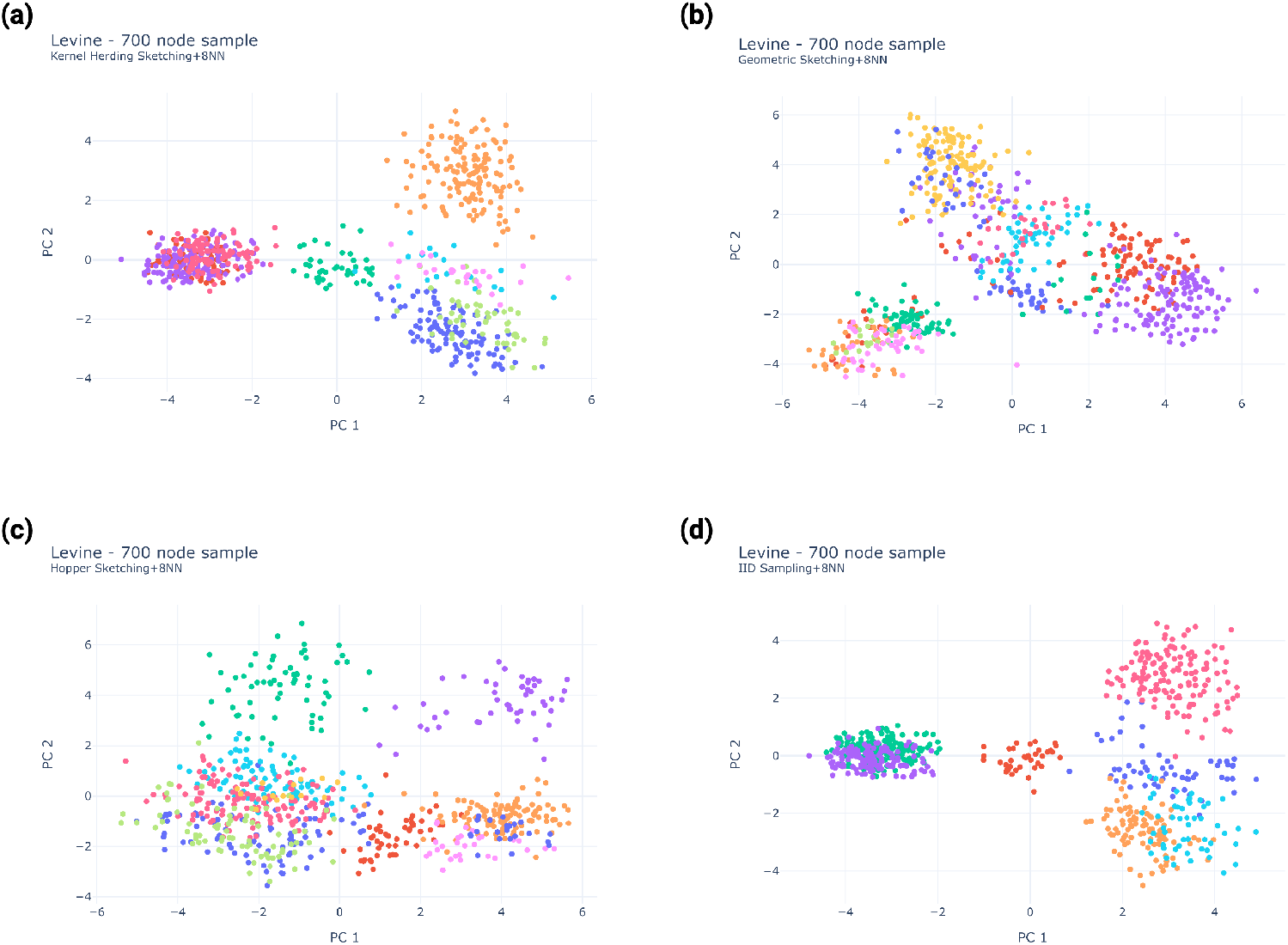
Samples of 700 cells from the Levine AML dataset projected in two dimensions with PCA and colored by Leiden clustering assignments. All sketching methods are shown here and include (a) Kernel Herding Sketching, (b) GeoSketching, (c) Hopper Sketching, and (d) IID sampling. A representative 8-NN graph was used in all visualizations. Regardless of sketching method, using an 8-NN graph significantly reduced the number of clusters produced by Leiden clustering. The Kernel Herding method (a) yields the most clear structure in the data.

**Figure 6:**
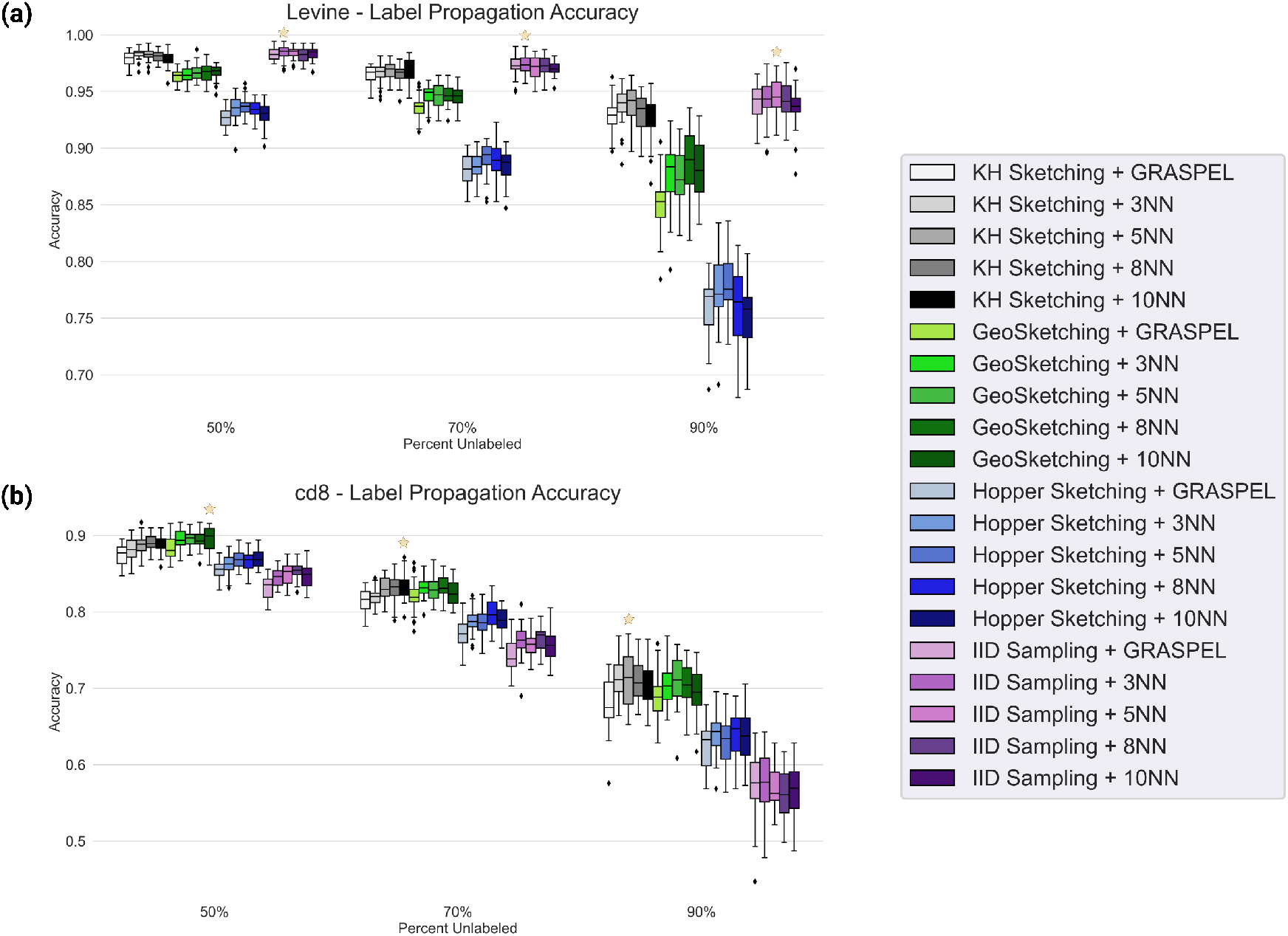
We formulated a semi-supervised label propagation task to evaluate the concordance between the graph structure and corresponding cell labels. Distributions over 30 sketches under each method are shown for (a) the Levine AML dataset and (b) the CD8 T-cell dataset. The star indicates the highest median accuracy for each number of nodes. We can see that graph generation method does not amplify or diminish performance, but downsampling method has a significant effect. In particular, the IID sampling and Kernel Herding methods perform best for the Levine dataset, and the Geometric Sketching and Kernel Herding Sketching methods have the highest accuracy on the CD8 T-cell dataset.

#### Kernel Herding

Kernel Herding Sketching has a single hyperparameter `gamma`, which controls the standard deviation of the distribution from which the random Fourier frequencies are selected.^1^ The standard deviation for the distribution is 1*/*`gamma`, so a high value of `gamma` leads to a tight distribution. We compared Kernel Herding sketches for `gamma`= 1, 5, 10, and 20 on population identification and semi-supervised learning, and we found that `gamma`=5 led to the best results.

#### Geometric Sketching

When sampling using Geometric Sketching, we used all of the default parameter values.^2^ In particular, we found that using the ‘`auto`’ option for specifying the number of covering boxes was optimal. This option sets the number of covering boxes as 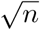, where *n* is the number of cells in the original data matrix. We also observed that varying the parameters `alpha` and `max_iter` from the default values of 0.1 and 200, respectively, did not significantly affect the downstream evaluations.

#### Hopper

Because each iteration of the Hopper algorithm depends on the previous one and the algorithm has to find an absolute maximizer every iteration, Hopper can become prohibitively time-consuming when working with large datasets. To speed up the process, the authors created the Treehopper method, which first partitions the data using principal components (where the maximum size of each subset is passed as a parameter) and then iterates on each section of the partition.^3^ Due to the size of the datasets used in these experiments, we used the Treehopper method with a PCA Tree Partition, where the maximum partition size was set at 1000 nodes. For all other parameters, we used the default values.

### 2.3 Implementation Details for Graph Generation Methods

Given that the GRASPEL algorithm uses a 2-NN graph as its base, we used its built-in k-NN graph implementation to generate the 3-NN, 5-NN, and 8-NN graphs.^4^ This implementation sets the edge weights of the edge *e*_*ij*_ to be

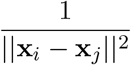

The GRASPEL algorithm takes in several hyperparameters: First, the `samplePerc` parameter determines the percentage of nodes at the top and bottom of the Fiedler vector to be considered for edge candidacy. Next, the `edge_iter` parameter determines how many edges should be added in each GRASPEL iteration, given as a percentage of the total nodes in the sample. The `sig` parameter provides the prior variance of the features. Finally, the `tol` and `maxIter` parameters provide stop conditions for the algorithm; when the spectral embedding distortion is below the value `tol` for all the candidate edges, the algorithm terminates. If this does not happen after `maxIter` iterations, then the algorithm terminates.

We used the default parameter values provided in the demo script: `samplePerc=1e-2, edge_iter=1e-4, sig=1000, tol=10,` and `maxIter=100`, respectively. We found that changing these parameter values did not significantly affect the downstream analysis.

## 3 Results

In order to best evaluate the utility of graph construction and downsampling methods for uncovering biologically cohesive cell-populations, we performed a range of experiments on two publicly available CyTOF and RNA-sequencing datasets. Individual cells in both datasets contain ground-truth cell labels, making evaluation of graph partitioning (e.g. identifying cohesive cell-populations) tractable. Experiments on all datasets were formulated to quantitatively explore the interplay between all sketching methods and graph generation strategies by partitioning the graphs under each particular combination with the Leiden clustering [15].

### Datasets

Here we describe the details of the two single-cell datasets used in all experiments.

### Levine AML Dataset

The Levine AML dataset is a CyTOF dataset with cells sampled from 16 patients with AML and 5 healthy controls [16] and accessed in Flow Repository under accession number FR-FCM-ZZPH. This resulted in 265,627 total cells with 32 measured protein features. The dataset has been classically used in benchmarking studies as a large subset of the cells have a manually gated ground-truth label. In our experiments, we restricted our view to these 104,184 labeled cells. The cells in this dataset are immune cells from patients with acute myeloid leukemia (AML). There are 14 distinct cell-type groups in this dataset; this is the label we used in our evaluation, since similar cell-types are often close to each other in feature space. Also, the cell-type groups are fairly well-separated in feature space, but the relative frequencies of the cell types vary widely; both of these factors affect the effectiveness of community detection.

### CD8 T-cell Dataset

The CD8 T-cell dataset [20] is a single-cell RNA sequencing dataset consisting of mouse CD8+ T cells profiled over 12-time points following infection with the Armstrong strain of the lymphocytic choriomeningitis virus (LCMV): Naive, d3-, d4-, d5-, d6-, d7-, d10-, d14-, d21-, d32-, d60-, d90-post infection. Single-cell RNA sequencing data were accessed from the Gene Expression Omnibus using the accession code GSE131847. The dataset was subsequently quality control filtered according to the distribution of molecular counts. To remove dead or dying cells, we filtered cells that had more than twenty percent of their total reads mapped to mitochondrial transcripts. Genes that were observed in less than three cells or had less than 400 counts were also removed. Following cell and gene filtering, the data were transcripts-per-million normalized and highly variable genes were selected using a normalized dispersion measure in scanpy v1.8.1 (`flavor=Seurat`, `min_mean=0.0125`, `max_mean=8`, `min disp=0.5`. The resulting dataset consisted of 45,207 cells and 2,539 gene features. We used principal component analysis (PCA) to reduce the dimension of this space to 30 principal components prior to downsampling. The main cell attribute that correlated with the data geometry was the sampling day (of which there were 12), so we used that as our label in the tests we ran.

### Cell Population Quality

First, we wanted to assess how well each method can recover ground-truth cell-populations through graph partitioning (e.g. graph-based clustering). In both datasets, individual cells had discrete labels determined by manual analysis (Levine AML dataset) or according to an experimental parameter (CD8 T-cell dataset). That is, in the Levine AML dataset, cells were labeled based on expert-annotated cell-populations, whereas in the CD8 T-cell dataset, cells were labeled according to their sampling day post LCMV infection. To test the capacity to identify cellular subsets in line with their ground-truth labels, each graph representation of the data was partitioned with the Leiden algorithm. Then, cell-to-cluster assignments were then evaluated based on the extent that they overlapped with the true labels with normalized mutual information (NMI). Geometric properties of the dataset, such as how well-separated clusters are, were quantified with the *Clustering Distance Ratio*, defined below.

First, normalized mutual information (NMI) was used to quantify the overlap between the inferred cell-to-cluster assignments and the ground-truth labels of cells. Briefly, NMI is a metric that quantifies how correlated two variables are and specifically quantifies what we can gather about one variable based on observing another. We compared the proposed clusterings from the Leiden algorithm to ground-truth labels for cells, which were cell-population in the Levine AML dataset and sampling day in the CD8 T-cell dataset. Possible values of NMI range from 0 to 1, where an NMI of 0 indicates entirely uncorrelated concordance between cellular ground-truth and cluststering, and an NMI of 1 indicates strong overlap. The results of computed NMIs quantifying concordance between true and inferred cellular labels across both datasets for all tested combinations of sketching and graph generation methods are shown in Figure 3.

Next, we designed a new metric called the clustering distance ratio (CDR) to quantify the extent of separation between clusters. Data with clear structure 1) have clusters which are separated in feature space and 2) contain dense sets of points within each cluster. Therefore, in defining the CDR, the numerator *n* is computed as the average pairwise Euclidean distance in feature space between the clusters, which addresses the first criterion. That is,

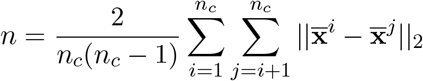

Here, *n*_*c*_ is the number of clusters, ||·||_1_ is the *L*_2_ (Euclidean) norm, and 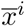 is the mean of the *i*^*th*^ cluster in feature space. The denominator of the CDR is further defined as the maximum cluster diameter, or the Euclidean distance between the two points that are farthest from each other within a cluster. Mathematically, this implies

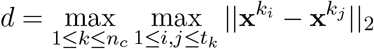

where *t*_*k*_ is the total number of nodes in the *k*^*th*^ cluster and 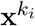 indexes the *i*^*th*^ node in the *k*^*th*^ cluster. We used Euclidean distance in both of these quantities so that the resulting ratio would be invariant under both orthogonal transformations and scalings. Because a large value of *n* indicates clusters that are well-separated from one another and a small value of *d* results from clusters where points are densely packed, this implies that a high CDR is more favorable. Results for the CDR across datasets, sketching methods, and graph generation metrics are shown in Figure 2.

To gain further qualitative intuition about the overall cluster structure of each combination of dataset, sketching algorithm, and graph generation technique, we visualized samples of 700 cells projected according to the first two principal components (Figs. 4 and 5). This allows us to qualitatively observe the cluster separation and to confirm that our metric CDR metric adequately captures our ideal notion of well-separated clusters.

### Semi-Supervised Learning to Quantify Overlap Between Cellular Labels and Graph Structure

Finally, we formulated a label propagation task to evaluate the overlap between node labels and graph structure. Briefly, label propagation (LP) is a semi-supervised learning algorithm which can infer the labels for unlabeled nodes in a partially-labeled dataset. In short, label propagation views the input graph as a discrete time Markov chain with labeled nodes as “sink” nodes (i.e. once a random walker encounters a labeled node, it terminates). The algorithm then calculates the probability *p*_*i,α*_ that a random walker starting at node *i* will encounter a node labeled *α* for all possible nodes and labels. These calculations can all be carried out simultaneously for one timestep via a matrix multiplication. The iteration stops either after a specified number of timesteps *t*_*max*_ or when the probabilities change by less than some specified tolerance *δ*_*max*_. In our experiments, we used *t*_*max*_ = 500 and *δ*_*max*_ = 0.001. Finally, each node is ultimately assigned the label with the highest probability, so that

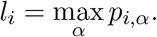

The accuracy of this process *acc*_*LP*_ was quantified as the percentage of labels guessed correctly by the LP algorithm as

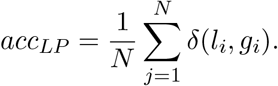

Here, *N* is the number of nodes in the sample, *δ* is the Kronecker delta function returning 1 if *l*_*i*_ = *g*_*i*_ and 0 otherwise, and **g** is the vector of ground-truth labels. High accuracy for the label propagation task implies more extensive overlap between the node labels and graph structure. We computed this accuracy metric for all combinations of sketching and graph generation strategies; those results are shown in Figure 6.

## 4 Discussion

Overall, our experiments inform principled strategies for uncovering biologically relevant cell-populations from single-cell data. First, our results revealed that the choice of sketching algorithm and the extent to which such sketches faithfully represent the overall data geometry are ultimately more important than how the graph is constructed. Although naive IID downsampling performed well in some tasks, the single-cell specific sketching methods exhibited comparable – or often stronger – performance in all scenarios. In particular, Geometric Sketching and Kernel Herding tended to be strong standouts across evaluations. From a graph construction point of view, *k*-NN graphs with *k* between 5 and 10 generally produced optimal results, with diminishing returns in performance with increasing graph density, but we also observed that the best choice of *k* was both dataset and task dependent. Overall, this suggests that classical *k*-nearest neighbor graphs built on quality sketches of the data uncover subsets of cells that are often higher quality than those discovered with sophisticated graph structure learning techniques, such as GRASPEL. In this section, we detail those experimental results and the intuition behind them.

Figures 2, 3, 4, and 5 broadly reflect the capacity of the Leiden clustering algorithm to uncover biologically meaningful sub-populations based on each sketching and graph generation combination. The boxplots in Figures 2 and 3 show the distribution of Leiden clustering quality results in terms of NMI and CDR metrics and can be further visually corroborated with two-dimensional visualizations shown in Figures 4 and 5. First, the CDR results in Figure 2(a) show that, when paired with a semi-dense graph (such as the 3NN or 5NN graph), Kernel Herding, Geometric Sketching, and IID sampling all perform pretty similarly to one another. We can verify this result qualitatively in Figures 4 and 5, where one sample from each sketching method is projected into two dimensions using PCA on the downsampled feature matrix. The clusters created on the 8-NN graph of the Hopper sketch (Fig. 5(a)) are much less clearly organized than in any of the other plots. Additionally, the shape of the data produced by the Hopper sketch appears to be markedly different from the other three.

Although Geometric Sketching does produce a high CDR on the Levine dataset, it underperforms on the CD8 T-cell dataset as compared to the Kernel Herding Sketching and IID sampling methods (Fig. 2(b)). The geometry of the data could be playing a role here; the groundtruth groups by sampling day are difficult to separate in feature space because of the continuity of the attribute of interest. The spatial covering technique utilized in the Geometric sketching method likely amplifies this difficulty. It is surprising to note that, with the exception of the ultra-sparse GRASPEL graph, the CDR generally decreases with increasing graph density in both datasets. This is possibly because the communities in this dataset are so distinct in feature space that the graph does not need many edges to capture those connections when the sketch is sufficiently good. Adding more low-confidence edges would add noise to the system, leading to lower-confidence clustering results.

The poor sketch quality of the Hopper method also affected the NMI results, as seen in Figure 3, where the Hopper method has significantly lower NMI results for all sample sizes. Interestingly, we see several differences in the data trends as compared with the CDR results of Figure 2. Firstly, Geometric Sketching has a poor performance with respect to the NMI metric, despite performing well under the CDR metric. This is likely because of the number of clusters prouced on the Geometric Sketching data. The groundtruth has 14 distinct cell types, which the Kernel Herding and IID sampling methods were able to capture efficiently. The Geometric Sketching method produced more clusters than the groundtruth cell types allow, indicating that the downsampling may have created some artificial separation within groundtruth clusters. Also, the NMI is also positively correlated with graph density on the Levine data; using 8-NN and 10-NN graphs produced the highest NMI within almost every sketching method group. This result is also affected in large part by the number of clusters produced by the Leiden algorithm. When there is less connection between nodes (as on sparse graphs), communities are less compressible; thus, the Leiden clustering algorithm produces more clusters. There are only 14 cell types in the Levine dataset, so a proposed clustering of 25 cell groups (as depicted in Fig 4 (b)) would likely not be highly correlated with the ground-truth. Because the Leiden algorithm prioritizes ensuring that the communities identified are well-connected, weakly connected communities (such as those produced by sparse graphs) will be incorrectly split by the algorithm into their better-connected subcommunities.

Note that there is a plateau in performance as graph density increases, with the elbow of this plateau starting at around 10 nearest neighbors. We ran our experiments on additional NN and 15-NN graphs (which were omitted for readability) and verified this plateau continued and almost entirely flattened out on these more connected graphs. The number of clusters produced via Leiden clustering stabilizes in this *k* range, and the additional edges added are essentially only contributing noise.

Interestingly, the reverse of the correlation seen in the Levine data is present in the CD8 T-cell dataset (Fig. 3(b)). Here, as graph density increases, the NMI generally decreases. One possible explanation for this difference is the nature of the labels we used. In the Levine dataset, we compared the Leiden clustering to the cell type groups, which is a discrete label. In the CD8 T-cell dataset, we used the sampling day, which is a continuous label; this makes the ground-truth cluster labels harder to cleanly identify. This implies that adding additional edges is more likely to introduce noise or spurious connections between the sampling day groups, thus making the ground-truth harder to recover. Furthermore, we observed that overly sparse graphs also hurt performance, as the ultra-sparse GRASPEL graph results often produced a lower NMI than that obtained using the 3-NN graph across almost all sketching groups. Also somewhat surprisingly, the Geometric Sketching method performs well on the CD8 T-cell data, a departure from the NMI results for the Levine data. In this dataset, the distribution of the cells is more even throughout the feature space, which means that the structure of the sketched data is more similar between sketching methods here than in the Levine dataset.

The last test we performed was label propagation to evaluate the general overlap between graph structure and node labels. Figure 6 displays label propagation accuracy as a function of varying percentages of unlabeled nodes. In this test on the Levine dataset, the IID sampling and Kernel Herding downsampling methods perform the best, with the IID sampling having a slightly higher accuracy. In agreement with the Leiden clustering task, the more weakly connected Hopper and Geometric Sketching methods struggle with label propagation. In both datasets, the graph generation method does not appear to have a consistent or significant effect on the accuracy. This is surprising, considering the apparent correlation between graph density and the other metrics. Related work by Wickramasinghe and Muthukumarana found in [31] that, in general, graph mining approaches like label propagation perform better on sparse graphs than they do on dense graphs, and graph density varies significantly from the ultra-sparse GRASPEL to the 8-NN graph. These results suggest that label propagation is more robust to noise and sparsity in the edges of a graph than Leiden clustering. In the CD8 T-cell dataset, Geometric Sketching and Kernel Herding consistently outperformed the other two sketching methods in the label propagation task.

While optimal choices of sketching and graph generation method are often task-dependent, there are several general takeaways from our experiments. Most importantly, a sketching method is generally advised over IID downsampling, with Kernel Herding and Geometric Sketching being the top performing methods across most experiments. Additionally, when the objective is to identify quality and biologically cohesive cellular subsets, a semi-dense *k*-NN graph is generally optimal. Finally, for semi-supervised learning tasks such as label propagation, the graph generation method used does not significantly affect performance, so it is best to use simple, efficient representations like *k*-NN graphs. In conclusion, our work suggests that quality sketching methods should generally be prioritized over graph generation strategy in order to identify high quality, biologically cohesive cellular subsets.

## Footnotes

1 https://github.com/CompCy-lab/SketchKH

2 https://github.com/brianhie/geosketch

3 https://github.com/bendemeo/hopper

4 https://github.com/Feng-Research/GRASPEL

